# Unlocking the Neuropeptidome using a Novel Endogenous Peptidomics Framework

**DOI:** 10.1101/2025.06.12.659356

**Authors:** Lauren Fields, Wenxin Wu, Tina C. Dang, Angel E. Ibarra, Mitchell Gray, Lingjun Li

**Affiliations:** Department of Chemistry, University of Wisconsin-Madison, 1101 University Avenue, Madison, WI 53706; School of Pharmacy, University of Wisconsin-Madison, 777 Highland Avenue, Madison, WI 53705

**Author notes:** Equal contribution (Lauren Fields, Wenxin Wu).

**Keywords:** Neuropeptide, EndoGenius, peptide, mass spectrometry, peptidomics, endogenous, spectral library, DIA, FAIMS

## Abstract

Endogenous peptides have garnered increasing attention over the past decade driven by the development of advanced analytical methods. However, large-scale investigations of peptides as potential disease biomarkers or drug candidates are still hindered by their challenging biochemical properties and the scarcity of specialized analytical tools. Among these, neuropeptides are particularly challenging to study due to their low *in vivo* concentration, rapid turnover rate, and high structural variability. Data-independent acquisition (DIA) mass spectrometry (MS) has shown great ability in profiling low-abundance ions. Nevertheless, most available DIA analytical tools are designed for proteomics studies and are not suitable for endogenous peptides, as there is no set enzymatic cleavage for these peptides. Here, we introduce the novel EndoGenius platform, paired with DIA-NN, to achieve high-confidence neuropeptide identification using an updated spectral library for DIA MS analysis. By employing orthogonal offline fractionation, ion mobility instrumentation, and an optimized database searching algorithm specifically for neuropeptides, we have constructed the largest crustacean neuropeptide spectral library to date. With this library, in combination with neural networking technology, we report a 100-fold increase in the number of neuropeptides identified in all *Cancer borealis* tissues analyzed. We also cross-validated these findings with transcriptomics data to enhance identification confidence. This workflow presents a novel analytical framework for DIA peptidomics analysis, offering a robust approach to studying neuropeptides and other endogenous peptides.

**TOC:** 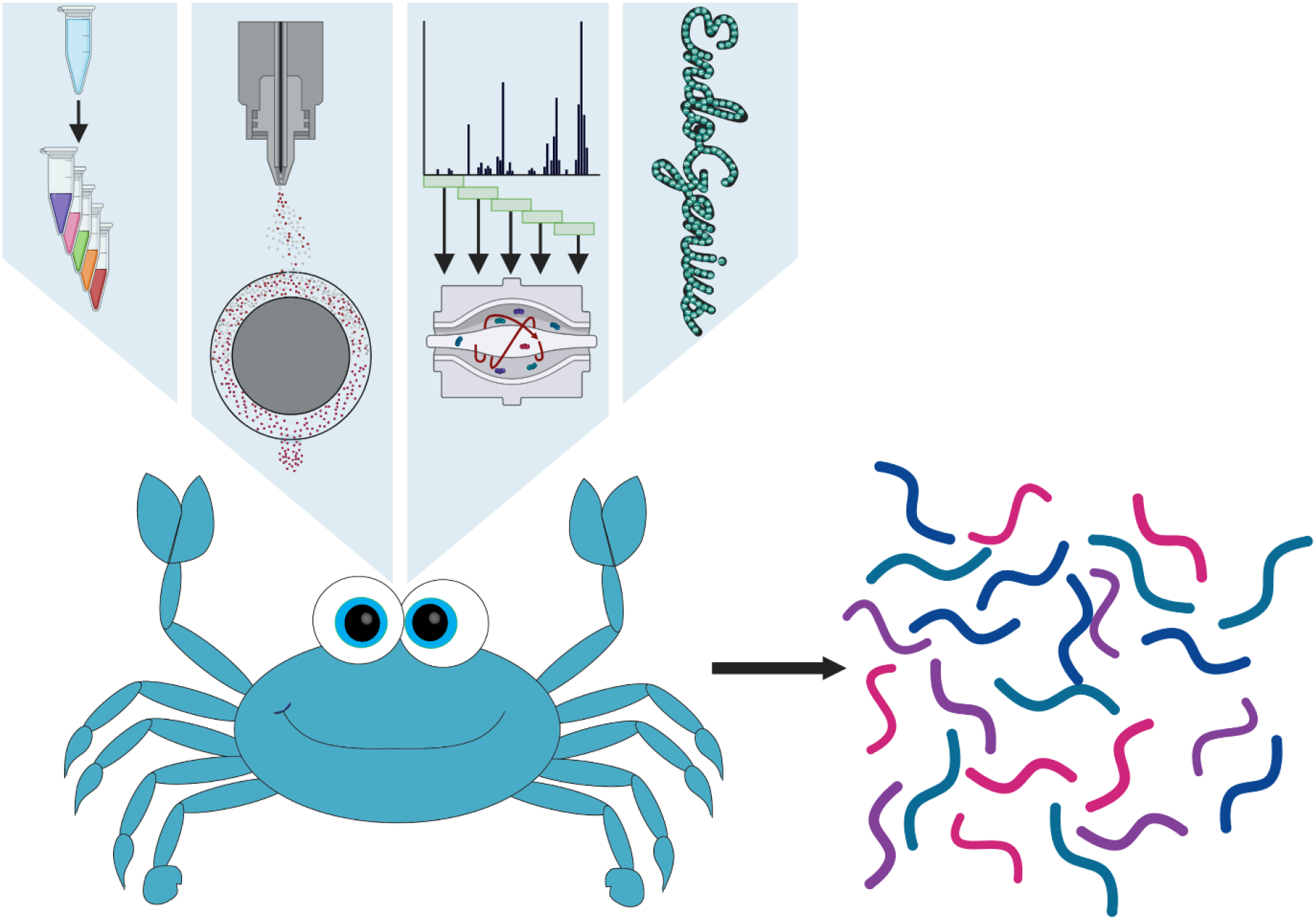

**Synopsis:** We present a framework that capitalizes on robust analytical innovations and an optimized bioinformatics pipeline to provide the most comprehensive snapshot of the crustacean neuropeptidome to-date.

## Introduction

Endogenous peptides are among the most informative molecules, responsible for numerous fundamental roles in all cellular domains^1^. Thus, much effort has been dedicated to determine the context in which these peptides are activated, understand their biological functions, and evaluate their potential as biomarkers for diseases or as key components in developing new treatments and medications. Clinically, more than 100 peptide are already approved by the Food and Drug Administration (FDA) for various applications, with many more in the discovery and clinical trial stages^2^.

Among endogenous peptides, neuropeptides are particularly important due to their significant therapeutic potential^3^. Neuropeptides are short chains of amino acids synthesized and released by neurons, acting as neuromodulators. They are cleaved from prohormones by endogenous proteases and released into circulating fluids or targeting tissues^4^. Neuropeptides serve as key signaling molecules in the nervous system, influencing biological processes such as food intake, mood regulation, and homeostasis maintenance^5-8^. They also play roles in many neurological dysfunctions, such as in Alzheimer’s disease, anxiety, and post-traumatic stress disorder^9-12^. Despite their significance as essential signaling molecules within the nervous system, neuropeptides are challenging to study due to many of their innate properties^13, 14^. Their fast turnover rate leads to rapid degradation, making them difficult to detect and analyze. Additionally, their low abundance *in vivo* often places them below the detection limits of many instruments^4^. Furthermore, their high structural complexity adds another layer of difficulty to their analysis, presenting significant challenges for researchers attempting to understand their functions and roles^3, 4^.

High-resolution mass spectrometry (HRMS) -based proteomics and peptidomics have emerged as invaluable tools for extensive peptide research^15^. Its ability to perform high-throughput profiling of biological samples, including those with low analyte concentration, makes HRMS peptidomic workflows particularly advantageous for both discovery-based studies and targeted analysis^16^. Traditionally, most MS methods are performed in data-dependent acquisition (DDA) mode. In DDA-MS, the top *n* most abundant precursor ions are selected for MS/MS fragmentation. Given that neuropeptides are typically present in especially low *in vivo* abundance (pM to fM)^17^, many neuropeptides remain undetected in DDA experiments. In contrast, data-independent acquisition (DIA) MS offers perhaps a more suitable alternative. In DIA-MS, all precursor ions are fragmented within predefined *m/z* windows^18^. While this provides a more comprehensive approach, subsequent data analysis is notoriously complicated, as often multiple precursor ions co-fragment in the same MS/MS spectrum^19, 20^. Despite this, DIA-MS provides a more comprehensive and unbiased dataset, with great potential for studying neuropeptides and their intricate roles in biological processes, albeit with additional consideration for data-analysis^21^.

Even with the development of advanced HRMS methods, studying neuropeptides remains challenging due to the lack of specialized software for analysis. Often, neuropeptide data analysis is conducted using platforms designed for proteomics applications, including MaxQuant^22^, PEAKS^23, 24^, and MSFragger^25^. While these tools have substantial value to their respectful fields, challenges are encountered when applying these for routine neuropeptidomic analyses. Lacking knowledge of endogenous cleavage patterns, neuropeptides are prepared for sample acquisition without enzymatic digestion^26^, a central component to bottom-up proteomics workflows. Further, neuropeptides have been identified in lengths ranging from three to over one hundred residues in length^4^. Additionally, distinct, active neuropeptides have been identified that differ in as little as a single residue^27^. Given this high level of sequence similarity, and neglect for the other stated considerations, we find that platforms developed for proteomics often over-identify a single neuropeptide. With this, we initially developed EndoGenius for optimized elucidation of neuropeptides from DDA-MS datasets^28^. However, strategies for the analysis of neuropeptides from DIA-based experiments are currently lacking.

The bioinformatics landscape with respect to DIA mass spectrometry has quickly expanded with many library-based and library-free options available^29, 30^. Spectral libraries traditionally represent a collection of high-resolution spectra with annotations for particular peptides^31^. These libraries can then be referenced to parse identifications from DIA fragmentation spectra. While these libraries are often very confident, they require additional samples and instrument time, making this a dedicated effort. Alternatively, library-prediction based tools such as Prosit^32^ and MS2PIP^33^ can be used to predict the spectra corresponding to a database which can be referenced to process DIA datasets. Lastly, platforms like DIA-Umpire work to deconvolute DIA spectra, generating pseudo-DDA spectra amenable to existing DDA platforms^34^. Previous studies have utilized DIA-Umpire for neuropeptide identification from DIA data collected from *Cancer borealis* neural tissues and circulating fluids^15, 35, 36^. While these searches confirmed the advantage of utilizing a DIA approach over a DDA, it has been noted that these deconvolution methods have inferior reproducibility compared to library-based methods^31, 35, 37^. A pilot study was recently conducted, in which a small spectral library was developed for *C. borealis*, which was produced from the accumulation of spectra from archived datasets. Despite the limited nature of this library, it revealed improved reproducibility of neuropeptide identifications from DIA datasets^35^.

Given the marked improvement of neuropeptide identification yielded by EndoGenius in the context of DDA experiments^28^, we sought to expand its application to DIA applications. By utilizing high-pH offline HPLC fractionation followed by online high-field asymmetric-waveform ion mobility spectrometry (FAIMS), a newer technology that affords an additional degree of separation, we report a spectral library that represents 58% percent of the known crustacean neuropeptidome. This is achieved from a bioinformatics perspective by utilizing EndoGenius to process all spectra. We then apply this library to parse DIA spectra obtained from *C. borealis* tissue. This was achieved through EndoGenius’ new spectral library building module, which is integrated with DIA-NN^38^, harnessing the power of neural network technology to produce the largest survey of the crustacean neuropeptidome to-date. Through this, we were able to illustrate the validation of neuropeptides that were previously predicted from transcriptomics, representing a milestone for neuropeptidomics^39-42^.

## Results and Discussion

### Considerations for Neuropeptide Spectral Libraries

Crustacea serve as a key model system for neuropeptidomic analyses, regarded for their simple nervous system that can be used to garner a deeper understanding of neuronal signaling. For example, *C. borealis*, has one of the most well-characterized central-pattern generating circuits in the animal kingdom for the pyloric rhythm and gastric mill behaviors, pertinent to feeding^43, 44^. Additionally, the neuroendocrine tissue of interest in this system are comprised of just a handful of neurons, enabling a fingerprint of neuropeptide activity to inform a deeper understanding of complex biological processes^45, 46^. Despite this, crustacean neuropeptides remain a challenging biomolecule to study, even with state-of-the-art MS instrumentation. This challenge is derived from the low *in vivo* abundance of neuropeptides, on a picomolar scale^47^, the rapid degradation of these molecules, and the reduced ionization of this species. This ionization issue is a result of the routine workflow for neuropeptide analysis, which does not include an enzymatic digestion step, typically used to improve ionization and reduce subsequent search space.

To alleviate these issues, we hypothesized that the generation of a spectral library for crustacean neuropeptides would enable the robust evaluation of these neuropeptides through DIA-MS. DIA-MS is particularly ideal for this task at hand, since in DIA, all precursor ions within a specified window, regardless of abundance undergo MS/MS fragmentation, a necessity for amino acid-level sequence resolution. Admittedly, this method produces far more complex output spectra, with multiple precursor ions simultaneously fragmented, compared to the more established method of data-dependent acquisition (DDA) MS. In DDA-MS, only one precursor ion at a time undergoes fragmentation, selecting ions for fragmentation on the basis of abundance, biasing against molecules of low abundance, such as neuropeptides. To address the complexities in the data analysis of DIA-MS spectra, spectral libraries are routinely used. Indeed, a pilot spectral library for crustacean neuropeptides was recently reported, showing promising results on a smaller scale^35^.

To build this spectral library where prioritization of resolution was kept in mind, we started with an abundance of neuroendocrine tissue (n=15) from the *C. borealis* pericardial organs (PO), brain, sinus glands (SG), thoracic ganglion (TG), and stomatogastric nervous system (STNS). Upon neuropeptide extraction, samples underwent fractionation using offline high-pH HPLC. Each fractionated sample underwent further separation using online HPLC before entering FAIMS and subsequently undergoing DDA MS/MS (**Figure 1**).

**Figure 1.**
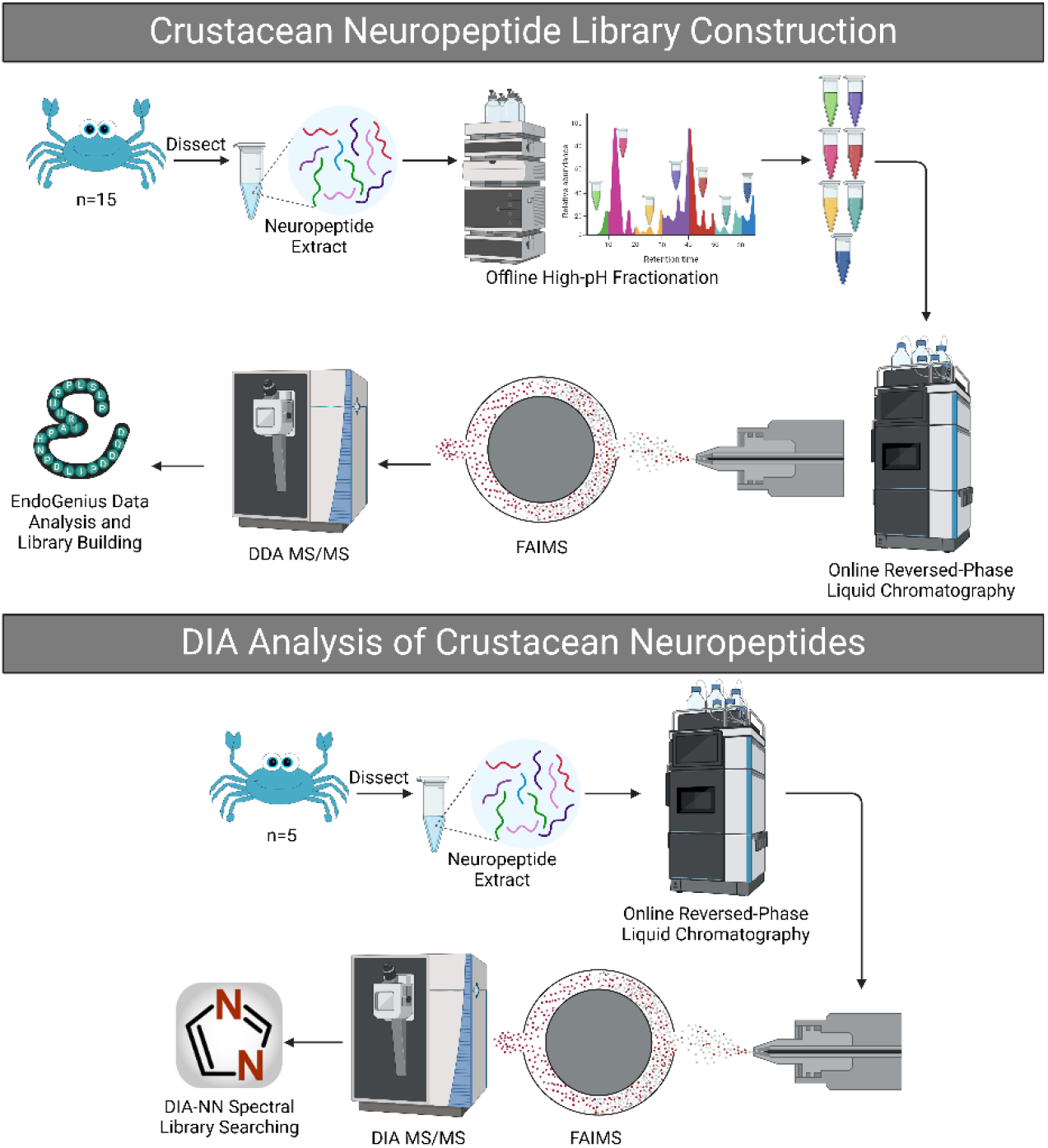
Overview of workflows used for construction of the spectral library using data-dependent mass spectrometry (DDA-MS) and subsequent analysis of data-independent acquisition (DIA) datasets. *C. borealis* crabs were first dissected, from which neuropeptides were extracted. Samples underwent offline, high-pH fractionation. Spectra were then acquired using online LC-FAIMS-MS/MS and processed using EndoGenius. Additional *C. borealis* tissue were dissected and extracted for DIA analysis, again acquired with online LC-FAIMS-MS/MS. The data were then searched against the spectral library using DIA-NN software.

### FAIMS and EndoGenius Expands Spectral Library

We hypothesized that the usage of FAIMS, a recently developed technology, would improve the performance of our dataset. FAIMS is a gas-phase-based ion separation method on the basis of ion mobility with respect to electric fields^48^. This method has been documented to decrease the interference of background ions, decreasing the limits of detection to improve the analysis of low-abundance analytes^48^. As neuropeptides are highly homologous and known to produce chimeric spectra, it was hypothesized that perhaps an additional measure of separation would benefit the offline and online HPLC methods already integrated in the workflow^4, 49^. To ensure greater separation, we introduced 3 compensation voltages (CV): -30 V, -45 V, and -60 V, using an internal stepping method. It should be noted that applying three CVs does contribute to an increased duty cycle, and that it is arguably more favorable in instances of abundant analytes to perform external stepping, using just a single CV at a time^50, 51^. However, these tradeoffs shift the balance to internal stepping in this scenario, where following fractionation, quantity of analyte is limited.

To further prioritize maintaining the most complete spectral library for crustacean neuropeptides documented to-date, we utilized EndoGenius for data analysis. EndoGenius is a software suite we recently presented optimized for the identification of neuropeptides^28^. This is achieved through utilizing conserved amino acid sequence motif regions to parse through sequences of high similarity, a common challenge with conventional mass spectrometry database searching platforms^27^. In addition, EndoGenius applies a scoring mechanism designed and optimized for non-tryptic peptides, further emphasizing its utility in this context. EndoGenius was used to perform database searching of all DDA spectra achieved through the fractionated LC-FAIMS-MS/MS data. From here, all spectra were merged strategically to construct a spectral library.

Duplicate entries, corresponding to unique spectra that each pertain to the same peptide, must be filtered from a complete spectral library. This can be done in two primary ways: scoring-based or cluster-based. In scoring-based library filtering, the peptide spectrum match (PSM) corresponding to the highest score is retained for the library. This approach is utilized in Bibliospec^52^, the library building component of Skyline^53, 54^, wherein the PSM with the highest dot product is retained. Another option is the cluster-based approach, wherein all PSMs corresponding to the same peptide are systematically merged to generate a single, composite spectrum. Software platforms including GLEAMS^55^ and MaRaCluster^56^ are able to achieve these merged spectra.

With a heightened appreciation that not all protein-derived calculations are directly amenable to neuropeptides, we sought to confirm which metric should be utilized to filter our spectral library for duplicate peptide entries. This evaluation was conducted on a series of criterion that are integrated into the EndoGenius scoring algorithm, including the average number of annotations per fragment ion, average fragment ions per amino acid, EndoGenius Score, hyperscore, motif score, number of fragment peaks, percent sequence coverage, and precursor intensity. Also included were normalized EndoGenius score, normalized hyperscore, and normalized precursor intensity, normalized with respect to the highest value within the library. It was evident that the normalized EndoGenius score and non-normalized hyperscore yielded more reproducible results across three technical replicates each of TG, Brain, and PO tissue types from the training dataset. Additionally, the hyperscore filter yielded a slight improvement in number of identifications, with respect to the normalized EndoGenius score (**Figure S1**). Given the established use of the hyperscore function^57^, originally implemented in X!Tandem^58^, for the correlation of neuropeptide mass spectra data, we elected to use the former, scoring-based approach, retaining the PSM with the highest hyperscore. We then built a spectral library building module addition for EndoGenius, available as a graphical-user interface (GUI) for increased accessibility (**Figure S2**).

### Evaluation of Spectral Library

We hypothesized that the combination of high-resolution instrumentation, a Thermo Vanquish Neo HPLC coupled to a Thermo Exploris 480 Orbitrap Mass Spectrometer, offline pre-fractionation of complex neuropeptide samples, and utilizing FAIMS technology and optimized bioinformatics data analysis would afford the most comprehensive crustacean neuropeptide spectral library to-date. We elected to first assess this hypothesis through benchmarking of the metrics of the spectral library described herein to the previously-published pilot spectral library, described recently^35^. It should be noted that the context of the pilot spectral library, from here referred to as the “original library” was that a library was constructed using an existing archive of non-fractionated DDA spectra, to illustrate the utility of using existing data to repurpose into a spectral library (**Supplemental File 1**). Thus, the library described herein, referred to as the “FAIMS library”, was generated with much more intention for the purpose of providing a complete picture of the neuropeptidome.

The overlap of the FAIMS library was first assessed, with the original library comprised of 239 unique peptide backbones, while the FAIMS library was comprised of 515 unique peptide backbones (**Figure 2A; Supplemental File 2**). Unique peptide backbones refer to the amino acid sequence alone, without consideration of any post-translational modifications (PTMs). This value reflects the diversity of the library and appreciates the library’s ability to parse through highly similar sequences. It should be noted that the known crustacean neuropeptidome is comprised of 891 unique peptide backbones, though this database includes entries predicted from transcriptomic datasets^4, 59, 60^ and from PEAKS *de novo* sequencing^21, 61^. Thus the original library conservatively represented approximately 26% of the crustacean neuropeptidome, while this work more than doubled the representation of the crustacean neuropeptidome to approximately 58%. We also observed that the distribution of peptide precursor charge was consistent between libraries (**Figure 2B**), in line with expected behavior for crustacean neuropeptides. Consideration of peptide length is important for the targeted spectral library herein, as crustacean neuropeptides in their non-tryptic form can have a length anywhere between three to one hundred amino acids^4, 62^. We see that in the original library, the peptides were identified with up to 56 residues in length, whereas in the FAIMS library, the longest peptide was just 38 residues in length (**Figure 2C**). There are a number of scenarios that this can be attributed to. For example, if the 60-residue peptide was low in concentration, it may have been lost during the separation steps. Perhaps most probable, the original library was analyzed using PEAKS Online. While in general we know that PEAKS does not afford as many neuropeptide identifications at a confident threshold as EndoGenius does, we do acknowledge PEAKS’ strength in its inclusion of *de novo* sequencing within its database searching algorithm^28^. Thus, it could be postulated that this particular neuropeptide was resolved with the aid of a joint *de novo* sequencing/database searching approach. The distribution of the five most common PTMs was consistent between both approaches (**Figure 2D**). The precursor *m/z* distribution was also similar between both libraries, matching the literature expectation that the majority of crustacean neuropeptides fall in the range of 400 to 800 *m/z* (**Figure 2E**)^21^. Based upon previously mentioned parameters and optimizations, we felt confident that we have compiled a spectral library that is fully representative of the known crustacean neuropeptidome.

**Figure 2.**
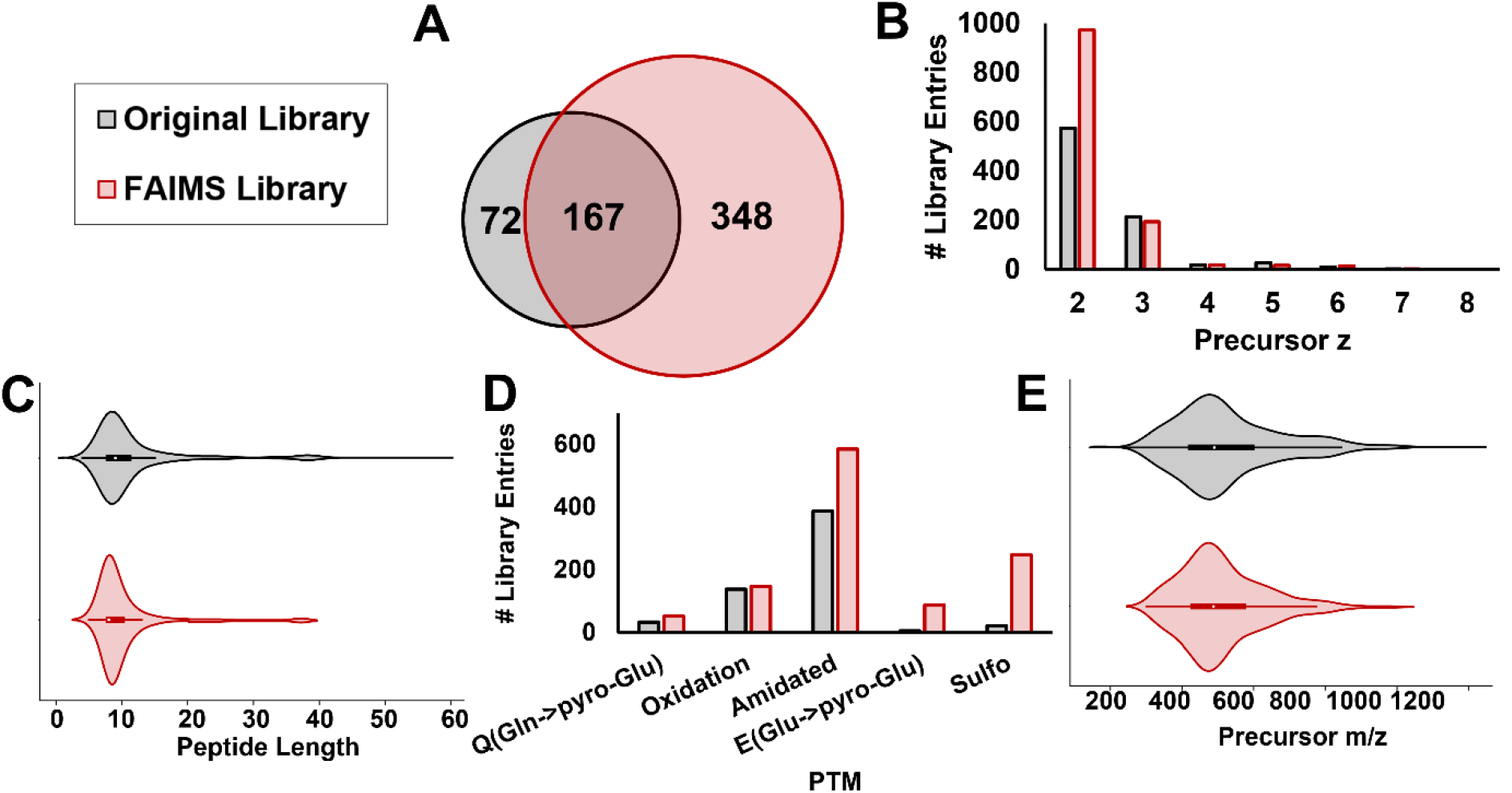
Comparison of library attributes from the original crustacean library to the library in this work. **(A)** This library incorporated more identifications while maintaining the same trends with regards to **(B)** precursor charge, **(C)** peptide length distribution, **(D)** post-translational modification frequency, and **(E)** precursor *m/z*.

We further sought to evaluate if these attributes described in Figure 2 were consistent across all samples included within the library. The samples include hemolymph (crustacean circulating fluid) and five key neuroendocrine tissue included with this study: brain, paired POs, paired SGs, STNS, and TG. This was imperative for us to ensure adequate representation of all neuropeptides for future behavioral studies. We found that the sequence length distribution (**Figure 3A**) and precursor retention time (**Figure 3B**) were consistent across all tissues. Finally, we determined the number of identifications yielded from each fraction through collection at 7-minute intervals. To ensure sufficient concentration of neuropeptides for analysis, fractions from less-abundant retention times, fractions 2-3 and 9-11 were pooled orthogonally. Indeed, we see that the number of unique IDs, inclusive of any PTMs, were contributed to by all tissues at all time points (**Figure 3C**).

**Figure 3.**
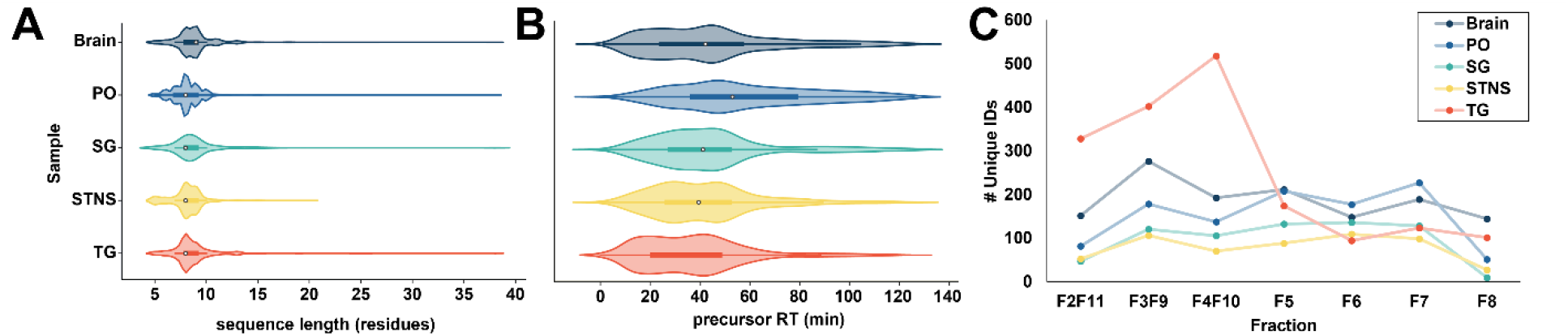
Survey of **(A)** sequence length and **(B)** precursor retention time (RT) for library entries attributes with respect to their tissue of origin. **(C)** number of unique IDs, inclusive of post-translational modifications, with respect to the tissue and fraction in which they were identified.

### DIA-NN Provides an Expanded Snapshot of the Neuropeptidome

To apply the improved neuropeptide spectral library for DIA analysis, we first collected data from the same respective tissues of 5 *C. borealis*. The sample preparation was consistent with those used for the spectral library generation, however, these samples did not undergo offline HPLC fractionation (**Figure 1**). This data was collected in DIA mode using LC-FAIMS-MS/MS. For data analysis, we turned to DIA-NN, a powerful platform for the interrogation of DIA mass spectrometry datasets^38^. This platform uses integrated neural networks for the evaluation of DIA-MS data in a time-efficient manner. Upon building our library in EndoGenius and formatting for DIA-NN, we seamlessly integrated DIA-NN into our platform. Upon doing so, we found a number of identifications across all 5 tissues examined that outperformed previous library-free^15, 21^ and library-based^35^ evaluations of the same sample types when considering unique peptide backbones (**Figure 4A; Supplemental File 3**) and unique identifications including PTMs (**Figure 4B**). While these results were promising without any prediction included on the part of DIA-NN, we further decided to evaluate the outcome of integrating this feature (**Supplemental File 4**). DIA-NN has a deep learning-based component that can predict spectra, retention time, and if applicable, ion mobility^38^. It should be noted that we used part of this feature, which builds upon a spectral library, but elected not to use the alternative prediction tool, which enables the digest of a FASTA file for library-free analysis. The feature was intentionally excluded so we could capitalize on our existing library yet use the predictive feature to integrate and inform new PTMs for searching. In this, we were able to identify approximately double the number of unique backbones between the standard DIA-NN and DIA-NN including prediction (**Figure 4A**). The power of this feature was far more apparent when evaluating the number of unique IDs, with up to a 100-fold increase in identified peptides (**Figure 4B**). This finding represents the most expansive characterization of the crustacean neuropeptidome to date. To gain an understanding of the unique IDs produced with the predictive feature, we compare the number of modifications present on each peptide and the identity of those PTMs. We found that most peptides identified with the predictive feature had three PTMs present, leading us to believe the strength of DIA-NN lies in the identification of extensively modified peptides (**Figure 4C**). These modifications included primarily C-terminal amidation, a ubiquitous modification across the crustacean neuropeptidome (**Figure 4D**). As neuropeptides often gain their functional roles through undergoing post-translational modifications, this work leads us to believe that we are able to utilize our spectral library to gain a greater understanding of biologically active, mature neuropeptides.

**Figure 4.**
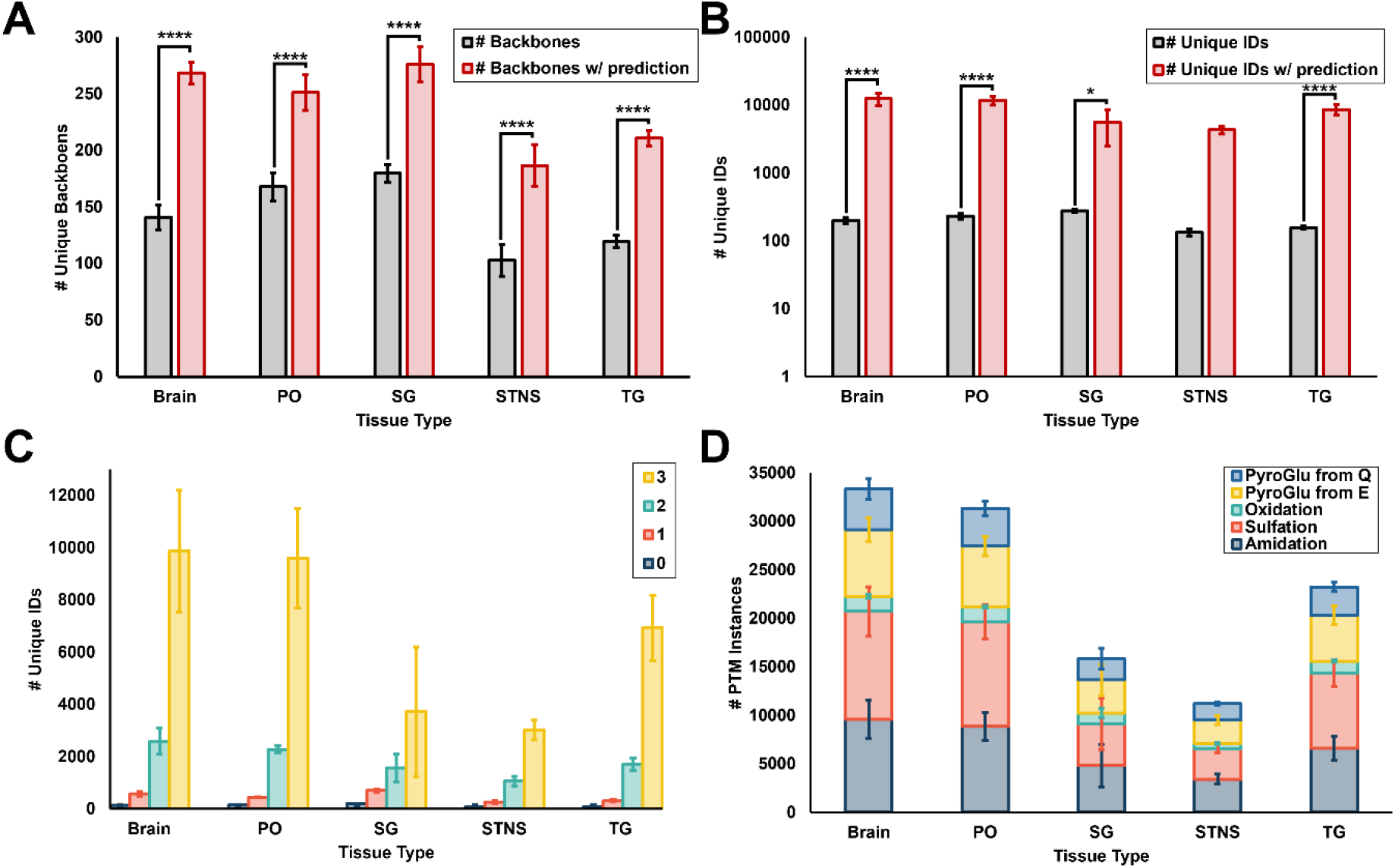
Results from search of DIA spectra obtained from neuropeptide extract of brain, PO, SG, STNS, and TG tissue types. Spectra were searched in DIA-NN using the spectral library generated in EndoGenius. Statistical comparison of results with and without the predictive feature were conducting using a two-way ANOVA with Bonferroni multiple comparisons test (* *p*-value < 0.05, ** *p*-value < 0.01, *** *p*-value <0.001,\*\*\*\**p*-value<0.0001). Results were searched with and without DIA-NN’s spectral prediction functionality, reported with the number of **(A)** unique IDs and **(B)** backbones, exclusive of any post-translational modification. Reported are **(C)** the number of PTMs identified per unique ID, with prediction, and **(D)** the identity of the PTM present on the unique ID. Values reported are average across three technical replicates with error bars representing standard deviation.

### DIA Remains Highly Reproducible

One of the primary advantages of DIA-MS is its superior reproducibility with respect to DDA-MS. In DDA, the precursors selected for fragmentation are selected on the basis of abundance, a stochastic approach that does not guarantee the repeated selection of the same precursor each time. While targeted methods such as parallel reaction monitoring (PRM) exist, these methods require an intimate understanding of the expected peptides within the sample. To address the biased nature of DDA, DIA instead fragments all precursors within a defined window size. Thus, reproducibility is superior in DIA^21^. Further, we have confirmed that reproducibility is increased when using a spectral library compared to depending on a spectral-library free deconvolution approach^35^, such as DIA-Umpire^34^, which seeks to match fragment ions with their respective precursor, essentially translating complex DIA spectra into a more manageable DDA format. Given the integration of this new platform, DIA-NN, we wanted to ensure that reproducibility of DIA spectra analysis was maintained. We also wanted to examine that the utilization of the spectra and retention time feature was not at the cost of reproducibility. When appreciating unique peptide backbones, we find a high level of reproducibility across three technical replicates within the spectra that did not include prediction (**Figure S3A-E**). Further, this trend was maintained, though on a larger scale, when the predictive feature of DIA-NN was included within the analysis (**Figure S3F-J**).

### Confirmation of Initial Transcriptomic Predictions

Traditional ways of studying neuropeptides were usually targeted, involving peptide purification and Edman degradation^63, 64^. Later on, RNA sequencing has been widely performed for larger scale molecular level investigation of the proteome level. Transcriptome prediction involves multiple steps of *in silico* enzymatic cleavages, increasing the potential diversity of the peptides identified^39^. Christie *et al*. in 2016 discovered 200 neuropeptides through *in silico* prediction from the neural transcriptome of *Cancer borealis*^41^. Using these predicted peptides together with MS identified *de novo* neuropeptides and other experimentally validated peptide sequences, we have built an in-house neuropeptide sequence library consisting of 892 peptide sequences (**Supplemental File 5**). In the identifications achieved from DIA-NN, most of the peptides identified were experimentally verified peptides. Additionally, we found that roughly 50 peptides from each tissue were identified for the first-time using MS, which were also predicted only from the transcriptome data^41^. For the *in silico* predicted peptides, there were also some peptides identified through other methods before transcriptome mining, and we have also identified them through MS (**Figure 5**). In this manner, our high-resolution MS results provide an additional layer of identification confidence for neuropeptides predicted *in silico* from the transcriptome (**Supplemental File 6**). The integration of transcriptomic and peptidomics data has significantly advanced our understanding of neuropeptide biology and their prohormone origin. Combining RNA sequencing with mass spectrometry can also allow us to correlate gene expression levels with peptide abundances, offering a comprehensive overview of neuropeptide regulation and function.

**Figure 5.**
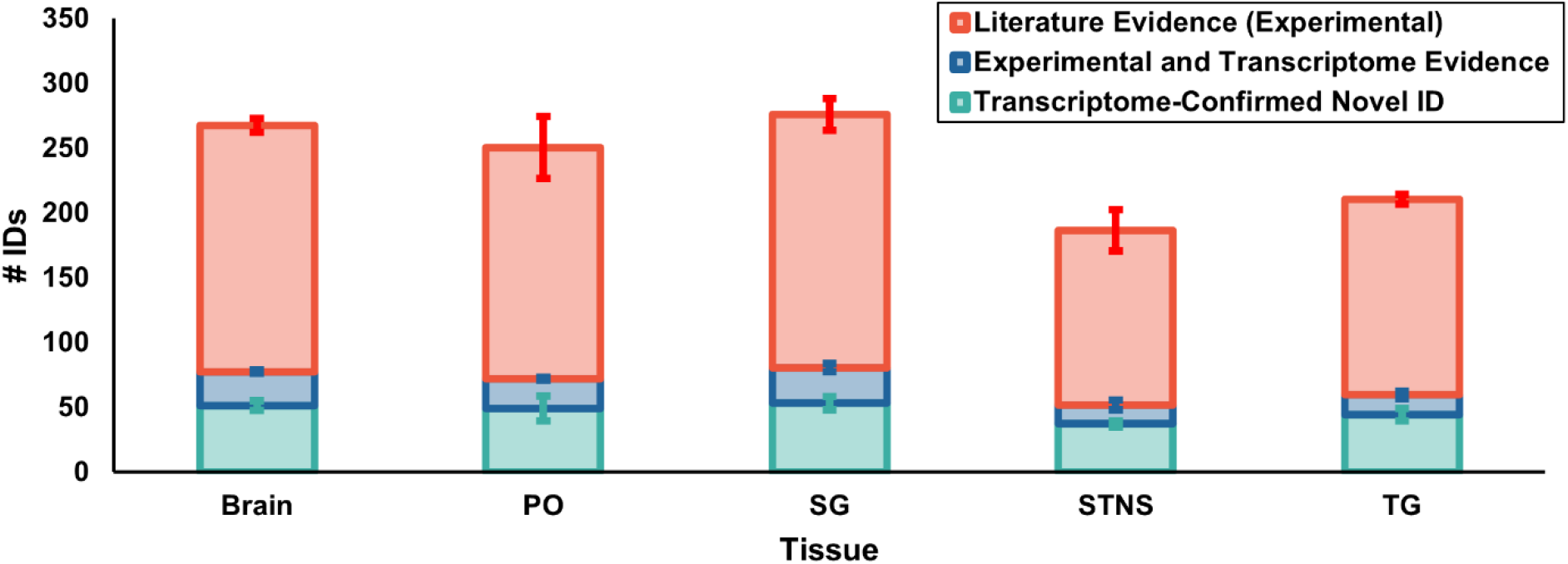
Identification achieved with spectra prediction from DIA-NN. Colors indicate different peptide origin. Most identified peptides are not from *in silico* transcriptome prediction but from experimentally-derived literature (red). Some peptides were identified in both previous literature and also transcriptome prediction (blue). Others were identified for the first time through mass spectrometry which was only predicted with in silico transcriptome mining (green). Values reported are the average across 3 technical replications with error bards representing standard deviation.

## Methods

### Materials and reagents

Optima grade formic acid was purchased from Sigma-Aldrich (St. Louis, MO). Desalting tips C18 ZipTip P10 were purchased from MilliporeSigma (Burlington, MA). The Stabilizor™ T1 was purchased from Denator AB (Gothenburg, Sweden). Methanol (MeOH), water (H_2_O), glacial acetic acid (GAA), sodium chloride (NaCl), potassium chloride (KCl), calcium chloride (CaCl_2_), magnesium chloride (MgCl_2_), sodium hydroxide (NaOH), and all other chemicals and solvents were purchased from Fisher Scientific (Pittsburgh, PA, USA). Sample preparation steps used ACS or HPLC grade solvent, and all mass spectrometry related steps used Optima grade solvents. Acidified MeOH was composed of 90/9/1 (v/v/v) MeOH/H_2_O/GAA. Crustacean saline solution was composed of 440 mM NaCl, 11 mM KCl, 13 mM CaCl_2_, 26 mM MgCl_2_, 11 mM Trizma® base, and 5 mM maleic acid, and the solution adjusted to pH 7.45 with NaOH.

### Animals and Dissection

Although no institutional approval is needed for working with crustacean, all experiments were performed following local or national regulations. Male *Cancer borealis*, were purchased from Global Market (Madison, WI) and were housed in artificial seawater tanks maintained at 10–12 °C, with a salinity of 35 parts per thousand (ppt) and 8-10 ppm of O_2_. The crabs were maintained on a 12 h light schedule exposure from 10 PM to 10 AM, and 12 h dark schedule with red light exposure from 10 AM to 10 PM daily. Crabs were allowed to adjust to the lab tank environment for at least 48 h prior to experiment. All crabs were anesthetized on ice for 20 min before sacrifice for collection of circulating fluid, hemolymph, and neuroendocrine tissues, as previously described^36, 65^. Neuropeptide rich tissues, including their brain, paired sinus glands (SG), paired pericardial organs (PO), thoracic ganglia (TG), and stomatogastric nervous system (STNS) were dissected in chilled crustacean saline solution.

### Sample preparation

Dissected tissues were heat stabilized using The Stabilizor™ T1 (Gothenburg, Sweden). After stabilization, tissues were flash frozen separately with dry ice. All procedures were performed on ice to prevent further degradation. 200 µL acidified MeOH per tissue was added to the samples before tissue homogenization. Tissues were homogenized with a hand-held ultrasonic homogenizer with 60 % amplitude, 8 s on, 15 s off until homogeneous. All samples were then centrifuged at 16,000 relative centrifugal force (RCF) for 30 minutes at 4 °C. The supernatant was dried down using speedvac on medium heat and stored at -80 °C until further use.

Hemolymph samples were extracted with 1:1 (v/v) of acidified MeOH, vortexed, bath sonicated and centrifuged at 16,000 RCF for 5 min at 4 °C. The supernatant was collected, and 0.5 equivalent of acidified MeOH was added to the pellet. The pellet was then manually homogenized with a pestle, vortexed, sonicated, and centrifuged again. This supernatant was collected, and the extraction process was repeated one more time. The supernatant was dried down using speedvac on medium heat and stored at -80 °C until further use.

### Off-line Fractionation

The dried samples were reconstituted in 508μL of Optima™ LC/MS Grade water with 10mM ammonium formate (pH=10). In total, five times of fractionations were done, with 100uL of sample fractionated each time (leaving 8μL dead volume). The samples were fractionated using a Kinetex® Core-Shell EVO 5um C18 column on a Waters Alliance 2695 Separations Module operating at 0.2mL/min with a column temperature at 30 °C. The mobile phase buffer A was 10mM ammonium formate, and mobile phase buffer B was 10mM ammonium formate with 90% ACN. The pH was adjusted to 10 using ammonium hydroxide for both phases. The samples were fractionated following a gradient as follows: 1% B to 35% B from 3 to 50 min, 35% B to 60% B from 50 to 54 min, and ramped to 70% B in 4 min, and the gradient was held at 70% B for 1 min and ramped to 100% B for 15 mins and ramped back to 1% B in 0.5mins and kept at 1% B for 19.5min for washing and equilibration. In total, 10 fractions from each 100 μL sample were collected for MS analysis, with each fraction containing 7 minute eluate from 7mins to 77mins. The first fraction (0-7mins) containing majorly salts was discarded. The collected fractions were then dried down with speedvac on medium heat and stored at −80 °C until further use.

### LC-MS/MS data acquisition

All Samples were reconstituted in water with 0.1% FA. Sample concentrations were tested using a NanoDrop™ 2000 (Thermo Fisher Scientific, CA) at a wavelength of 205 nm and equal amount of peptide sample were loaded onto a 15 cm length, 75 μm i.d. in-house-packed Bridged Ethylene Hybrid C18 (1.7 μm, 130 Å) column and were analyzed on an Orbitrap Exploris™ 480 Mass Spectrometer (Thermo Fisher Scientific, CA) coupled to a Vanquish Neo UPLC system (Thermo Fisher Scientific, CA). Mobile phase A was water with 0.1% FA, and mobile phase B was 80% acetonitrile (ACN) with 0.1% FA. Fractionated samples were analyzed in DDA mode with one technical replication and another set of the samples were used for DIA library validation with three technical replications using the following LC and MS parameters. Both DDA and DIA were analyzed using the same LC gradient and similar MS parameters to ensure reproducibility. Peptide samples were separated in online low-pH RPLC at a flow rate of 0.3 µL/min with a 126 min gradient (excluding the pre-and post-equilibration: 3% to 40% phase B from 0 min to 8 min, then 40% to 60% phase B in 25 min, the gradient then ramped to 90% phase B in 0.5min and held at 90% for 9.5 min followed by another ramp to 95% phase B in 0.5 min; the gradient was then held at 95% phase B for 10 min followed by column equilibration. For MS acquisition, samples were analyzed in positive ion mode with 2100 V static voltage, ion transfer tube temperature at 305 °C, 50% RF lens value and a static total carrier gas flow at 4.2 L/min. Three FAIMS compensation voltages (CV) were set for both DDA and DIA experiments, using an internal stepping method: -45V, -60V, and -75V. For both DDA and DIA, full MS scans were acquired at *m/z* 300–1500 with a resolving power of 60,000 and AGC target of 1E6 with a maximum injection time of 150 ms. A 1.5 s scan time was used for DDA MS/MS data acquisition. MS/MS acquisition was at a resolving power of 15,000 and AGC target of 1E5 with a maximum injection time of 250 ms. Only peaks with a charge state of 2-8 were fragmented through higher-energy collision dissociation (HCD) with a normalized collision energy (NCE) of 30%. Automatic detection for dynamic exclusion was used. For DIA, all other MS/MS parameters were the same as DDA MS/MS method except the cycle time was set to be 3 s and a 10 *m/z* isolation window with 1 *m/z* window overlap was used.

### Data analysis

All DDA spectra were processed in EndoGenius, as described previously^28^. All analyses were conducted in EndoGenius version 1.0.9. Prior to database searching, Thermo .RAW files were converter to .MS2 and .mzML formats in RawConverter^66^ and MSConvert^67^, respectively. Analysis parameters consisted of: parent mass error tolerance, 20 ppm; fragment mass error tolerance, 0.02 Da; m/z range, 50-3000 m/z; minimum precursor intensity, 1000; maximum precursor charge, 8; maximum fragment charge, 4; maximum post-translational modifications (PTMs) per peptide, 3; confident coverage threshold percent, 70%; EndoGenius Score threshold, 1000. PTMs included C-terminal amidation, oxidation of methionine, N-termini cyclization of glutamic acid and glutamine, and sulfation of tyrosine. These particular modifications were selected as they are well-documented within crustacean neuropeptide literature^68, 69^.

### DIA-NN Method Optimization

Previously, we established a method for searching spectral libraries in PEAKS Online, searching against our original spectral library. With this, we had knowledge of the expected outcomes of this dataset and used this for training our method established herein. Training data included the brain, PO, and TG crustacean tissues, available online^35^. These data were then searched in DIA-NN against the original spectral library (**Supplemental File 1**). We first selected parameters for DIA-NN utilization that were intuitive for neuropeptide work. These included: number of threads, 4; verbose, 1; Q-value, 0.01; variable modifications, 3; protein-group level, 1; minimum peptide length, 3; interfering precursor removal, 0. Output quantities matrices, reanalyse, heuristic protein inference, smart profiling, global mass calibration, global normalization, isoleucine and leucine equivalent reannotation, no SwissProt preference, and peak translation functionalities were applied. PTMS included those applied to DDA methods. To ensure a digest-free search was applied, cuts at all amino acids were disabled. To streamline analysis using the above parameters, DIA-NN was operated via the command line.

To build a spectral library from the aforementioned fractionated dataset, we first concatenated all results of all DDA EndoGenius searches. It is standard practice to ensure only one spectral library entry is present per peptide. To achieve this, we generated eleven cursory spectral libraries, each filtered on a separate criterion. Criterion included the average number of annotations per fragment ion, average number of fragment peaks per amino acid, EndoGenius score, hyperscore, motif score, number of fragment peaks present for the peptide, percent sequence coverage, and precursor intensity. Additional criterion were normalized EndoGenius score, normalized hyperscore, and normalized precursor intensity. Normalized calculations were performed relative to the highest respective value within the concatenated series of peptide-spectrum matches (PSMs). Each search was conducted in DIA-NN following the stated parameters and evaluated for reproducibility and number of yielded identifications.

Upon determination of the most effective library filtering manner, all FAIMS-DIA data were searched in DIA-NN. The final spectral library is available in **Supplemental File 5**. Prior to searching, spectra were converted to .mzML format, using MSConvert^67^. The spectral library was provided in .TSV format. All other parameters were consistent with those above.

### EndoGenius Spectral Library Module

A module was added to EndoGenius v.2.0, which includes functionality to build a spectral library from EndoGenius output reports. Provided with the directory where EndoGenius output reports are stored, the module searches all reports and concatenates into a large list of PSMs, as described above. The user also provides a fragment error threshold, to determine which peaks are eligible for library inclusion. The module then retrieves output reports and corresponding raw spectra, to obtain the retention time for the peptide. Duplicate PSMs are then removed, retaining the PSM with the highest hyperscore. The library is then formatted in a .TSV form, compatible with DIA-NN. This library includes the precursor m/z, product m/z, retention time, library intensity, modified peptide sequence, precursor charge, fragment series number, fragment type (b or y), fragment loss type, and fragment charge.

## Conclusions

The work presented herein illustrates the improved peptidomic depth that can be achieved through the combination of a comprehensive spectral library, high-resolution data-independent acquisition mass spectrometry, and optimized bioinformatics methodology. This work revealed a greater than 20-fold increase in neuropeptide identifications across all tissues, an achievement that will undoubtably propel the elucidation of neurochemical signaling mechanisms within crustacea. Further, we report the confirmation of neuropeptides that were previously only predicted from transcriptomics datasets, bolstering support for utilizing this approach to expand the neuropeptidome. Finally, as novel crustacean neuropeptides are identified, this dataset can be re-mined and the library expanded. This framework also highlights the opportunity to use innovative analytical and bioinformatics methods to analyze discrete biomolecules that are traditional challenging to identify, with the potential to apply to other translational models of interest.

## Data Availability

EndoGenius is open-source and freely available from GitHub at https://github.com/lingjunli-research/EndoGenius-v2.0. All mass spectrometry proteomics data have been deposited to the ProteomeXchange Consortium via the MassIVE partner repository with the dataset identifier MSV000095447.

## Acknowledgements

This work was supported in part by National Science Foundation (CHE-2108223) and National Institutes of Health (NIH) through grant R01DK071801. The Orbitrap instruments were purchased through the support of an NIH shared instrument grant (S10RR029531) and Office of the Vice Chancellor for Research and Graduate Education at the University of Wisconsin-Madison. LF was supported in part by the National Institute of General Medical Sciences of the National Institutes of Health under Award Number T32GM008505 (Chemistry–Biology Interface Training Program), the 2024 Eli Lilly and Company/ACS Analytical Graduate Fellowship, and a predoctoral fellowship supported by the NIH, under Ruth L. Kirschstein National Research Service Award (NRSA) from the National Institutes of Health-General Medical Sciences F31GM156104. T.C.D. was supported in part by the National Institute of General Medical Services of the National Institute of Health under Award 5T32GM141013 (Molecular and Cellular Pharmacology Training Program) and SciMed Graduate Research Scholars Fellowship through the University of Wisconsin-Madison. A.E.I. was supported in part by the National Science Foundation Graduate Research Fellowship Program under Grant No. DGE-2137424. Support was also provided by the Graduate School, part of the Office of Vice Chancellor for Research and Graduate Education at the University of Wisconsin-Madison, with funding from the Wisconsin Alumni Research Foundation and the UW-Madison. L.L. would like to acknowledge NIH grants RF1AG052324, R01AG078794, R21AG065728, S10OD028473, and S10OD025084, as well as funding support from a Vilas Distinguished Achievement Professorship and Charles Melbourne Johnson Professorship with funding provided by the Wisconsin Alumni Research Foundation and University of Wisconsin-Madison School of Pharmacy. The content is solely the responsibility of the authors and does not necessarily represent the official views of the National Institutes of Health or National Science Foundation. Figure 1 was generated using BioRender.

## References

1. Davenport, A. P.; Scully, C. C. G.; de Graaf, C.; Brown, A. J. H.; Maguire, J. J. Advances in therapeutic peptides targeting G protein-coupled receptors. Nat. Rev. Drug Discov. 2020, 19, 389–413.

2. Wang, L.; Wang, N.; Zhang, W.; Cheng, X.; Yan, Z.; Shao, G.; Wang, X.; Wang, R.; Fu, C. Therapeutic peptides: current applications and future directions. Signal Transduct. Target. Ther. 2022, 7 (1), 48.

3. Hokfelt, T.; Broberger, C.; Xu, Z. Q.; Sergeyev, V.; Ubink, R.; Diez, M. Neuropeptides— an overview. Neuropharmacology 2000, 39 (8), 1337–1356.

4. Christie, A. E.; Stemmler, E. A.; Dickinson, P. S. Crustacean neuropeptides. Cell Mol. Life Sci. 2010, 67 (24), 4135–4169.

5. Chen, R.; Hui, L.; Cape, S. S.; Wang, J.; Li, L. Comparative neuropeptidomic analysis of food intake via a multifaceted mass spectrometric approach. ACS Chem. Neurosci. 2010, 1 (3), 204–214.

6. An, S.; Harang, R.; Meeker, K.; Granados-Fuentes, D.; Tsai, C. A.; Mazuski, C.; Kim, J.; Doyle, F. J., 3rd; Petzold, L. R.; Herzog, E. D. A neuropeptide speeds circadian entrainment by reducing intercellular synchrony. Proc. Natl. Acad. Sci. 2013, 110 (46), E4355–4361.

7. Lin, S.; Boey, D.; Herzog, H. NPY and Y receptors: lessons from transgenic and knockout models. Neuropeptides 2004, 38 (4), 189–200.

8. Vu, J. P.; Larauche, M.; Flores, M.; Luong, L.; Norris, J.; Oh, S.; Liang, L. J.; Waschek, J.; Pisegna, J. R.; Germano, P. M. Regulation of Appetite, Body Composition, and Metabolic Hormones by Vasoactive Intestinal Polypeptide (VIP). J. Mol. Neurosci. 2015, 56 (2), 377–387.

9. Podvin, S.; Jiang, Z.; Boyarko, B.; Rossitto, L. A.; O’Donoghue, A.; Rissman, R. A.; Hook, V. Dysregulation of Neuropeptide and Tau Peptide Signatures in Human Alzheimer’s Disease Brain. ACS Chem. Neurosci. 2022, 13 (13), 1992–2005.

10. Schmeltzer, S. N.; Herman, J. P.; Sah, R. Neuropeptide Y (NPY) and posttraumatic stress disorder (PTSD): A translational update. Exp. Neurol. 2016, 284 (Pt B), 196–210.

11. Primeaux, S. D.; Wilson, S. P.; Cusick, M. C.; York, D. A.; Wilson, M. A. Effects of altered amygdalar neuropeptide Y expression on anxiety-related behaviors. Neuropsychopharmacology 2005, 30 (9), 1589–1597.

12. Wu, W.; Ma, M.; Ibarra, A. E.; Lu, G.; Bakshi, V. P.; Li, L. Global neuropeptidome profiling in response to predator stress in rat: Implications for post-traumatic stress disorder. J. Am. Soc. Mass Spectrom. 2023, 34 (8), 1549–1558.

13. Romanova, E. V.; Sweedler, J. V. Peptidomics for the discovery and characterization of neuropeptides and hormones. Trends Pharmacol. Sci. 2015, 36 (9), 579–586.

14. Stemmler, E. A.; Bruns, E. A.; Cashman, C. R.; Dickinson, P. S.; Christie, A. E. Molecular and mass spectral identification of the broadly conserved decapod crustacean neuropeptide pQIRYHQCYFNPISCF: the first PISCF-allatostatin (Manduca sexta-or C-type allatostatin) from a non-insect. Gen. Comp. Endocrinol. 2010, 165 (1), 1–10.

15. Phetsanthad, A.; Carr, A. V.; Fields, L.; Li, L. Definitive Screening Designs to Optimize Library-Free DIA-MS Identification and Quantification of Neuropeptides. J. Proteome Res. 2023, 22 (5), 1510–1519.

16. Hellinger, R.; Sigurdsson, A.; Wu, W.; Romanova, E. V.; Li, L.; Sweedler, J. V.; Süssmuth, R. D.; Gruber, C. W. Peptidomics. Nat. Rev. Methods Primers. 2023, 3 (1), 25.

17. Sauer, C. S.; Li, L. Multiplexed quantitative neuropeptidomics via DiLeu isobaric tagging. In Methods in Enzymology, Vol. 663; Academic Press Inc., 2022; pp 235–257.

18. Hu, A.; Noble, W. S.; Wolf-Yadlin, A. Technical advances in proteomics: new developments in data-independent acquisition. F1000Res 2016, 5, 419–419.

19. Heil, L. R.; Fondrie, W. E.; McGann, C. D.; Federation, A. J.; Noble, W. S.; MacCoss, M. J.; Keich, U. Building Spectral Libraries from Narrow-Window Data-Independent Acquisition Mass Spectrometry Data. J. Proteome Res. 2022, 21 (6), 1382–1391.

20. Pino, L. K.; Just, S. C.; MacCoss, M. J.; Searle, B. C. Acquiring and Analyzing Data Independent Acquisition Proteomics Experiments without Spectrum Libraries. Mol. Cell. Proteomics 2020, 19 (7), 1088–1103.

21. DeLaney, K.; Li, L. Data Independent Acquisition Mass Spectrometry Method for Improved Neuropeptidomic Coverage in Crustacean Neural Tissue Extracts. Anal. Chem. 2019, 91 (8), 5150–5158.

22. Cox, J.; Mann, M. MaxQuant enables high peptide identification rates, individualized p.p.b.-range mass accuracies and proteome-wide protein quantification. Nat. Biotechnol. 2008, 26 (12), 1367–1372.

23. Ma, B.; Zhang, K.; Hendrie, C.; Liang, C.; Li, M.; Doherty-Kirby, A.; Lajoie, G. PEAKS: powerful software for peptide de novo sequencing by tandem mass spectrometry. Rapid Commun. Mass Spectrom. 2003, 17 (20), 2337–2342.

24. Zhang, J.; Xin, L.; Shan, B.; Chen, W.; Xie, M.; Yuen, D.; Zhang, W.; Zhang, Z.; Lajoie, G. A.; Ma, B. PEAKS DB: de novo sequencing assisted database search for sensitive and accurate peptide identification. Mol. Cell. Proteomics 2012, 11 (4), M111 010587.

25. Kong, A. T.; Leprevost, F. V.; Avtonomov, D. M.; Mellacheruvu, D.; Nesvizhskii, A. I. MSFragger: ultrafast and comprehensive peptide identification in mass spectrometry-based proteomics. Nat. Methods 2017, 14 (5), 513–520.

26. Southey, B. R.; Amare, A.; Zimmerman, T. A.; Rodriguez-Zas, S. L.; Sweedler, J. V. NeuroPred: a tool to predict cleavage sites in neuropeptide precursors and provide the masses of the resulting peptides. Nucleic Acids Res. 2006, 34.

27. Vu, N. Q.; Yen, H. C.; Fields, L.; Cao, W.; Li, L. HyPep: An Open-Source Software for Identification and Discovery of Neuropeptides Using Sequence Homology Search. J. Proteome Res. 2023, 22 (2), 420–431.

28. Fields, L.; Vu, N. Q.; Dang, T. C.; Yen, H. C.; Ma, M.; Wu, W.; Gray, M.; Li, L. EndoGenius: Optimized Neuropeptide Identification from Mass Spectrometry Datasets. J. Proteome Res. 2024.

29. Shahbazy, M.; Ramarathinam, S. H.; Illing, P. T.; Jappe, E. C.; Faridi, P.; Croft, N. P.; Purcell, A. W. Benchmarking Bioinformatics Pipelines in Data-Independent Acquisition Mass Spectrometry for Immunopeptidomics. Mol. Cell Proteomics 2023, 22 (4), 100515.

30. Lou, R.; Cao, Y.; Li, S.; Lang, X.; Li, Y.; Zhang, Y.; Shui, W. Benchmarking commonly used software suites and analysis workflows for DIA proteomics and phosphoproteomics. Nat. Commun. 2023, 14 (1), 94.

31. Barkovits, K.; Pacharra, S.; Pfeiffer, K.; Steinbach, S.; Eisenacher, M.; Marcus, K.; Uszkoreit, J. Reproducibility, Specificity and Accuracy of Relative Quantification Using Spectral Library-based Data-independent Acquisition. Mol. Cell. Proteomics 2020, 19 (1), 181–197.

32. Gessulat, S.; Schmidt, T.; Zolg, D. P.; Samaras, P.; Schnatbaum, K.; Zerweck, J.; Knaute, T.; Rechenberger, J.; Delanghe, B.; Huhmer, A.; et al. Prosit: proteome-wide prediction of peptide tandem mass spectra by deep learning. Nat. Methods 2019, 16 (6), 509–518.

33. Degroeve, S.; Maddelein, D.; Martens, L. MS2PIP prediction server: compute and visualize MS2 peak intensity predictions for CID and HCD fragmentation. Nucleic Acids Res. 2015, 43 (W1), W326–330.

34. Tsou, C. C.; Avtonomov, D.; Larsen, B.; Tucholska, M.; Choi, H.; Gingras, A. C.; Nesvizhskii, A. I. DIA-Umpire: comprehensive computational framework for data-independent acquisition proteomics. Nat. Methods 2015, 12 (3), 258–264, 257 p following 264.

35. Fields, L.; Ma, M.; DeLaney, K.; Phetsanthad, A.; Li, L. A crustacean neuropeptide spectral library for data-independent acquisition (DIA) mass spectrometry applications. Proteomics 2024, e2300285.

36. DeLaney, K.; Hu, M.; Wu, W.; Nusbaum, M. P.; Li, L. Mass spectrometry profiling and quantitation of changes in circulating hormones secreted over time in Cancer borealis hemolymph due to feeding behavior. Anal. Bioanal. Chem. 2022, 414 (1), 533–543.

37. Zhang, F.; Ge, W.; Huang, L.; Li, D.; Liu, L.; Dong, Z.; Xu, L.; Ding, X.; Zhang, C.; Sun, Y.; et al. A Comparative Analysis of Data Analysis Tools for Data-Independent Acquisition Mass Spectrometry. Mol. Cell. Proteomics 2023, 22 (9), 100623.

38. Demichev, V.; Messner, C. B.; Vernardis, S. I.; Lilley, K. S.; Ralser, M. DIA-NN: neural networks and interference correction enable deep proteome coverage in high throughput. Nat. Methods 2020, 17 (1), 41–44.

39. Ma, M.; Gard, A. L.; Xiang, F.; Wang, J.; Davoodian, N.; Lenz, P. H.; Malecha, S. R.; Christie, A. E.; Li, L. Combining in silico transcriptome mining and biological mass spectrometry for neuropeptide discovery in the Pacific white shrimp Litopenaeus vannamei. Peptides 2010, 31 (1), 27–43.

40. Christie, A. E.; Cashman, C. R.; Brennan, H. R.; Ma, M.; Sousa, G. L.; Li, L.; Stemmler, E. A.; Dickinson, P. S.; Christie, A. E.; Cashman, C. R.; et al. Identification of putative crustacean neuropeptides using in silico analyses of publicly accessible expressed sequence tags. Gen. Comp. Endocrinol. 2008, 2 (156).

41. Christie, A. E.; Pascual, M. G. Peptidergic signaling in the crab Cancer borealis: tapping the power of transcriptomics for neuropeptidome expansion. Gen. Comp. Endocrinol. 2016, 237, 53–67.

42. Christie, A. E.; Lundquist, C. T.; Nässel, D. R.; Nusbaum, M. P. Two novel tachykinin-related peptides from the nervous system of the crab Cancer borealis. J. Exp. Biol. 1997, 200 (17), 2279–2294.

43. Cook, A. P.; Nusbaum, M. P. Feeding state-dependent modulation of feeding-related motor patterns. J. Neurophysiol. 2021, 126 (6), 1903–1924.

44. Nusbaum, M. P.; Beenhakker, M. P. A small-systems approach to motor pattern generation. Nature 2002, 417 (6886), 343–350.

45. Christie, A. E.; Skiebe, P.; Marder, E. Matrix of neuromodulators in neurosecretory structures of the crab Cancer borealis. J. Exp. Biol. 1995, 198 (Pt 12), 2431–2439.

46. Skiebe, P. Neuropeptides are ubiquitous chemical mediators: Using the stomatogastric nervous system as a model system. J. Exp. Biol. 2001, 204 (Pt 12), 2035–2048.

47. Sauer, C. S.; Li, L. Mass Spectrometric Profiling of Neuropeptides in Response to Copper Toxicity via Isobaric Tagging. Chem. Res. Toxicol. 2021, 34 (5), 1329–1336.

48. Saba, J.; Bonneil, E.; Pomiès, C.; Eng, K.; Thibault, P. Enhanced Sensitivity in Proteomics Experiments Using FAIMS Coupled with a Hybrid Linear Ion Trap/Orbitrap Mass Spectrometer. J. Prot. Res. 2009, 8 (7), 3355–3366.

49. Batth, T. S.; Francavilla, C.; Olsen, J. V. Off-line high-pH reversed-phase fractionation for in-depth phosphoproteomics. J. Prot. Res. 2014, 13 (12), 6176–6186.

50. Cooper, H. J. To What Extent is FAIMS Beneficial in the Analysis of Proteins? J. Am. Soc. Mass Spectrom. 2016, 27 (4), 566–577.

51. Creese, A. J.; Shimwell, N. J.; Larkins, K. P.; Heath, J. K.; Cooper, H. J. Probing the complementarity of FAIMS and strong cation exchange chromatography in shotgun proteomics. J. Am. Soc. Mass Spectrom. 2013, 24 (3), 431–443.

52. Frewen, B.; MacCoss, M. J. Using BiblioSpec for creating and searching tandem MS peptide libraries. Curr Protoc Bioinformatics 2007, Chapter 13 (1), 13 17 11–13 17 12.

53. MacLean, B.; Tomazela, D. M.; Shulman, N.; Chambers, M.; Finney, G. L.; Frewen, B.; Kern, R.; Tabb, D. L.; Liebler, D. C.; MacCoss, M. J. Skyline: an open source document editor for creating and analyzing targeted proteomics experiments. Bioinformatics 2010, 26 (7), 966–968.

54. Pino, L. K.; Searle, B. C.; Bollinger, J. G.; Nunn, B.; MacLean, B.; MacCoss, M. J. The Skyline ecosystem: Informatics for quantitative mass spectrometry proteomics. In Mass Spectrometry Reviews, John Wiley and Sons Inc.: 2020; Vol. 39, pp 229–244.

55. Bittremieux, W.; May, D. H.; Bilmes, J.; Noble, W. S. A learned embedding for efficient joint analysis of millions of mass spectra. Nat. Methods 2022, 19 (6), 675–678.

56. The, M.; Kall, L. MaRaCluster: A Fragment Rarity Metric for Clustering Fragment Spectra in Shotgun Proteomics. J. Proteome Res. 2016, 15 (3), 713–720.

57. Craig, R.; Beavis, R. C. A method for reducing the time required to match protein sequences with tandem mass spectra. Rapid Commun. Mass Spectrom. 2003, 17 (20), 2310–2316.

58. Craig, R.; Beavis, R. C. TANDEM: matching proteins with tandem mass spectra. Bioinformatics 2004, 20 (9), 1466–1467.

59. Christie, A. E. Identification of putative neuropeptidergic signaling systems in the spiny lobster, Panulirus argus. Invert Neurosci 2020, 20 (1), 2.

60. Tu, S.; Xu, R.; Wang, M.; Xie, X.; Bao, C.; Zhu, D. Identification and characterization of expression profiles of neuropeptides and their GPCRs in the swimming crab, Portunus trituberculatus. PeerJ 2021, 9, e12179.

61. DeLaney, K.; Hu, M.; Hellenbrand, T.; Dickinson, P. S.; Nusbaum, M. P.; Li, L. Mass Spectrometry Quantification, Localization, and Discovery of Feeding-Related Neuropeptides in Cancer borealis. ACS Chem. Neurosci. 2021, 12 (4), 782–798.

62. Keller, R. Crustacean neuropeptides: structures, functions and comparative aspects. Experientia 1992, 48 (5), 439–448.

63. Hetru, C.; Li, K. W.; Bulet, P.; Lagueux, M.; Hoffmann, J. A. Isolation and structural characterization of an insulin‐related molecule, a predominant neuropeptide from Locusta migratoria. Eur. J. Biochem. 1991, 201 (2), 495–499.

64. Tatemoto, K. Neuropeptide Y: complete amino acid sequence of the brain peptide. Proc. Nat. Acad. Sci. 1982, 79 (18), 5485–5489.

65. Gutierrez, G. J.; Grashow, R. G. Cancer borealis stomatogastric nervous system dissection. JoVE 2009, (25), e1207.

66. He, L.; Diedrich, J.; Chu, Y. Y.; Yates, J. R., 3rd. Extracting Accurate Precursor Information for Tandem Mass Spectra by RawConverter. Anal. Chem. 2015, 87 (22), 11361–11367.

67. Chambers, M. C.; Maclean, B.; Burke, R.; Amodei, D.; Ruderman, D. L.; Neumann, S.; Gatto, L.; Fischer, B.; Pratt, B.; Egertson, J.; et al. A cross-platform toolkit for mass spectrometry and proteomics. Nat. Biotechnol. 2012, 30 (10), 918–920.

68. Anapindi, K. D. B.; Romanova, E. V.; Checco, J. W.; Sweedler, J. V. Mass Spectrometry Approaches Empowering Neuropeptide Discovery and Therapeutics. Pharmacol. Rev. 2022, 74 (3), 662–679.

69. Akhtar, M. N.; Southey, B. R.; Andren, P. E.; Sweedler, J. V.; Rodriguez-Zas, S. L. Evaluation of database search programs for accurate detection of neuropeptides in tandem mass spectrometry experiments. J. Proteome Res. 2012, 11 (12), 6044–6055.

