## Supporting Information for "Unlocking the Neuropeptidome using a Novel Endogenous Peptidomics Framework"

[*]Corresponding author

**Table of Contents**

Supplemental Figures (located within this document)


- **Figure S1:** Spectral library filtering
- **Figure S2:** EndoGenius spectral library building GUI
- **Figure S3:** Reproducibility of DIA-NN results generated with and without prediction


Supplemental Files


- **Supplemental File 1:** Original spectral library
- **Supplemental File 2:** FAIMS and original spectral library overview
- **Supplemental File 3:** Identifications from search without prediction
- **Supplemental File 4:** Identifications from predictive search
- **Supplemental File 5:** FAIMS spectral library
- **Supplemental File 6:** Survey of neuropeptide accession origins

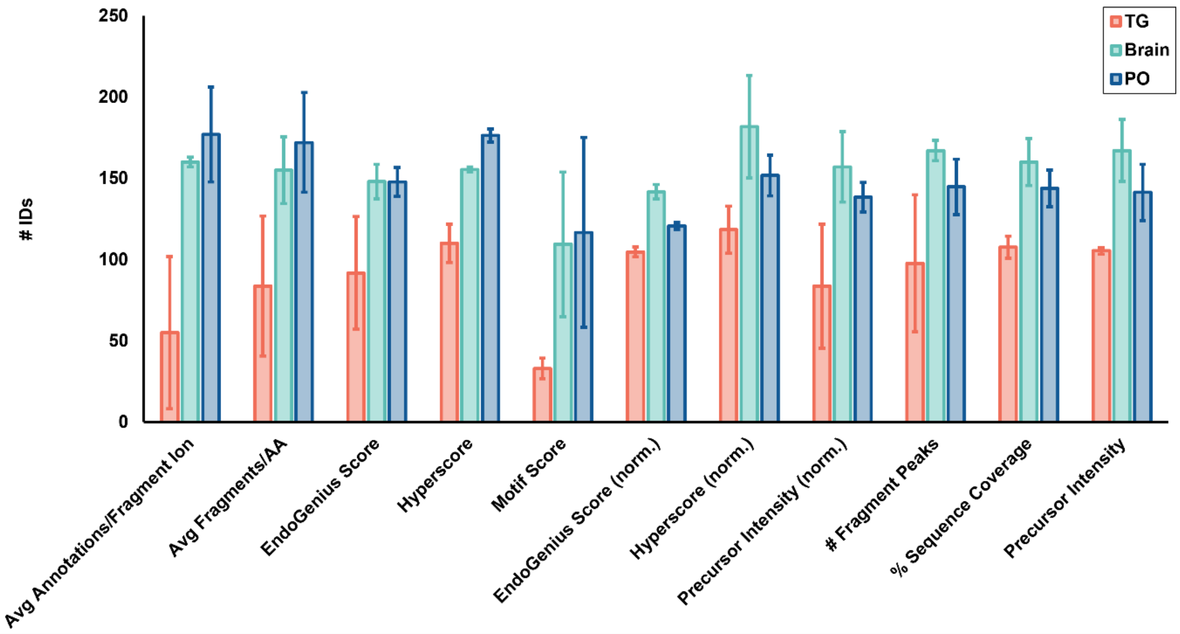


**Figure S1:** Number of identifications resulting from searching eleven spectral libraries, each filtered in accordance with a unique PSM metric. Bars represent the average ± the standard deviation of 3 technical replicates. Norm. indicates that all values were normalized to the highest value within the spectral file for the indicated metric.

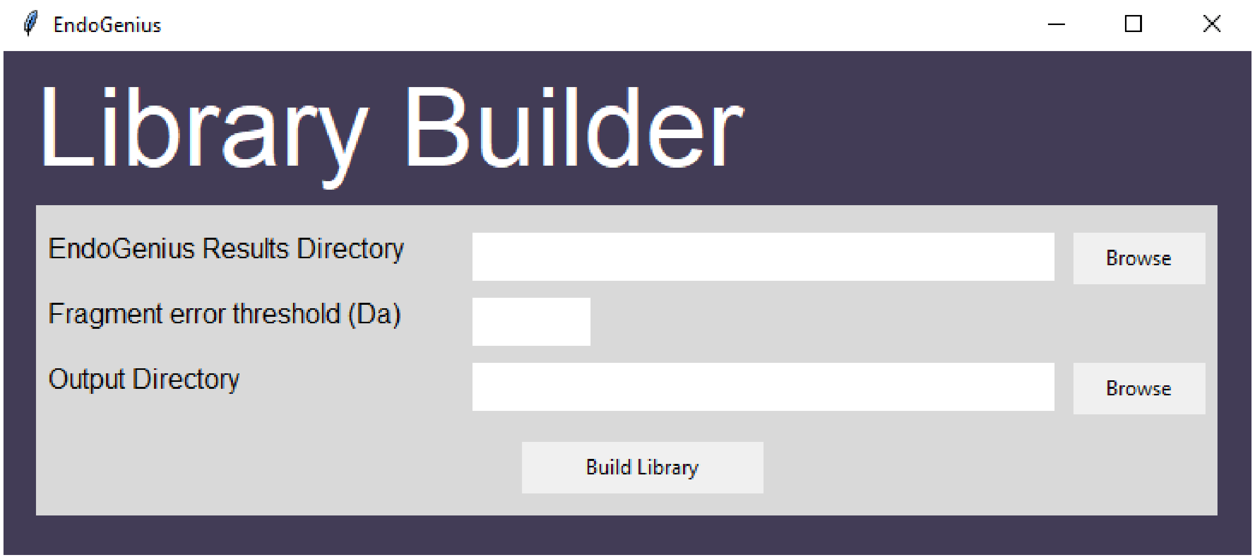


**Figure S2:** Library building module was constructed and implemented into EndoGenius. Users can upload EndoGenius results files, which will be mined to build a spectral library compatible with DIA-NN.

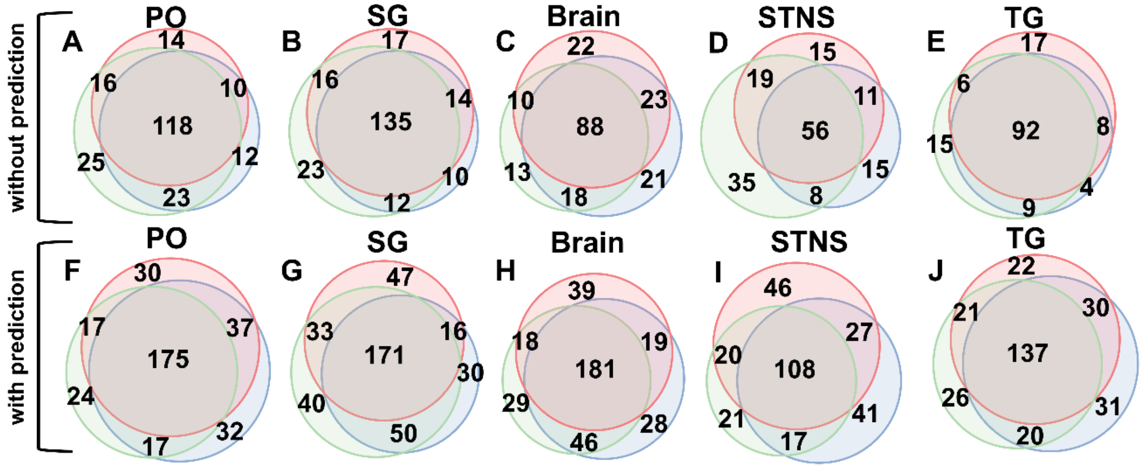


**Figure S3:** Overlap of identifications from three technical replicates produced by DIA analysis. Technical replicates 1, 2, and 3 are outlined in red, green, and blue, respectively. **(A-E)** Representation of unique backbones, limited to amino acid sequence alone, identified without prediction in DIA-NN. **(F-J)** Representation of unique backbones identified utilizing the predictive feature in DIA-NN.
